# GENVISAGE: Rapid Identification of Discriminative and Explainable Feature Pairs for Genomic Analysis

**DOI:** 10.1101/2020.02.05.935411

**Authors:** Silu Huang, Charles Blatti, Saurabh Sinha, Aditya Parameswaran

## Abstract

**Motivation:** A common but critical task in genomic data analysis is finding features that *separate* and thereby help explain differences between two classes of biological objects, e.g., genes that explain the differences between healthy and diseased patients. As lower-cost, high-throughput experimental methods greatly increase the number of samples that are assayed as objects for analysis, computational methods are needed to quickly provide insights into high-dimensional datasets with tens of thousands of objects and features.

**Results:** We develop an interactive exploration tool called Genvisage that rapidly discovers the most discriminative feature pairs that best separate two classes in a dataset, and displays the corresponding visualizations. Since quickly finding top feature pairs is computationally challenging, especially when the numbers of objects and features are large, we propose a suite of optimizations to make Genvisage more responsive and demonstrate that our optimizations lead to a *400X* speedup over competitive baselines for multiple biological data sets. With this speedup, Genvisage enables the exploration of more large-scale datasets and alternate hypotheses in an interactive and interpretable fashion. We apply Genvisage to uncover pairs of genes whose transcriptomic responses significantly discriminate treatments of several chemotherapy drugs.

**Availability:** Free webserver at http://genvisage.knoweng.org:443/ with source code at https://github.com/KnowEnG/Genvisage

## 1 Introduction

A common approach to discovery in biology is to construct experiments or analyses that directly contrast two specific classes of biological objects. Examples of this include examining patient samples contrasting tumor versus normal tissue [1], studying the differences in molecular effects of two competing drug treatments [2], or characterizing differentially expressed genes versus genes with unaltered gene expression in a carefully designed experiment [3]. To understand the mechanisms that determine these object classes, researchers often employ statistical and machine learning tools to identify a manageable subset of features, e.g. genes, that accentuate, discriminate, or help explain the differences between classes, i.e., *separate* the two classes. We therefore refer to this problem as the *separability problem*.

### Challenge 1: Explainable Separation with Guarantees

Many tools have been developed in several different biological settings [4–9] that attempt to solve the separability problem by focusing on discovering pairs of features that taken together strongly discriminate the classes. Feature pair methods can provide a better characterization of what distinguishes two object classes by offering insights into the interplay of important features that would not be found using single feature statistical tests [10] or univariate classifiers [11]. Specifically, predictors built with gene feature pairs are more robust to normalization and can achieve better model performance than predictors using single genes as features [5, 6]. On the other hand, methods focused on feature pairs offer the advantage of providing more interpretable or explainable results over more complicated machine learning approaches that return a complex combination of several features to discriminate the classes, such as multivariate regression with LASSO regularization [12] or pattern mining from random forest models [13]. Some existing papers [7, 14–16] employ these more complex machine learning approaches to heuristically return more interpretable feature pairs. However, these heuristic methods do not fully explore the search space nor do they offer a guarantee on the quality of the returned feature pairs.

### Challenge 2: Scalability in Data Size

A major problem with current methods that address the separability problem with either feature pairs or more complex machine learning models is that they do not scale to the growing size of genomic data sets. As is often the case with genomics, the biological objects being analyzed (e.g., tissue samples or drug experiments) are frequently represented by high dimensional numeric feature vectors (e.g., transcript abundance measurements). Additionally, with the rise of low-cost sequencing, the possible number of biological objects in a dataset is also increasing and likely to grow in orders of magnitude over the next decade [17]. Applying the standard methods to datasets with tens of thousands of objects and features results in massive running times that preclude interactive exploration of the data. For example, exhaustively searching for the optimal feature pairs from the full space of possibilities in a typical genomic analysis resulted in running times over an hour on a 200 node compute cluster in Watkinson et al. [9].

One reason that more complex machine learning and feature pair based methods do not scale well with the number of features and objects is the the selection of the metric for scoring separability. In Watkinson et al. [9], a metric called synergy is proposed for evaluating the utility of feature pairs, aiming to capture both linear and non-linear aspects of the separability of the two class. Consequently, the intrinsic complexity of these metrics makes them difficult to benefit from optimization techniques. Metrics based only on quantifying *linear* separability, on the other hand, may return a more limited subset of interesting features, but they also may be more intuitive for users to understand and simultaneously enable more performance optimizations and speedups. The linear separability metric has been used in previous studies to identify pairs of genes with expression differences between two cancer types [18] or pairs of motifs that discriminate between different types of genomic sequences [19].

### Our Proposal Genvisage: A Scalable and Explainable Tool for Addressing Separability

Motivated by these observations, we present GENVISAGE, an interactive data exploration tool designed to address the separability problem and scale to the size of large genomic analysis datasets. With GENVISAGE, we not only achieve high separability quality with our carefully formulated problem; but also enable explanations on separation via intuitive visualization; meanwhile, we are capable to handle large scale datasets efficiently — the best of all three world. Specifically, to enable this scalability, GENVISAGE focuses on returning the top ranking feature pairs that discriminate the objects of separate classes, rather than returning larger subsets of features using more complex and longer to train machine learning approaches. Genvisage is also based around a linear separability metric that provides an intuitive interpretation to feature pairs while enabling and simplifying the design of several important performance optimizations. These optimizations include *(a)* elimination of repeated computation for different features pairs; *(b)* pruning poor ranking pairs during early execution; *(c)* sampling with a quality guarantee to further reduce running time; and *(d)* cleverly traversing the search space of feature pairs for improved efficiency.

We applied Genvisage to two large genomic datasets with tens of thousands of objects and high-dimensional feature vectors where it is computationally expensive to score the separability for all possible feature pairs. In one, called LINCS, we find pairs of genes whose expression discriminates between perturbagen experiments involving different drug treatments, and in the other, called MSigDB, we find pairs of annotations (such as pathway membership) that separate differentially expressed cancer genes from other genes. With the carefully designed separability metric of Genvisage and its suite of sophisticated optimizations that accelerates evaluation, we are able to *accurately return the highest ranking separating feature pairs for both datasets within two minutes on a single machine*. This reflects a *180X* and *400X* speedup over a competitive baseline for the MSigDB and LINCS data sets (respectively). We also show that the feature pairs identified by Genvisage often more significantly discriminate between the object classes than the corresponding best ranking individual features, even after accounting for the larger search space. Finally, we performed an in-depth analysis for nine distinct drug treatments in the LINCS dataset and found 1070 feature (gene) pairs that had significant separability scores. These gene pairs were enriched in literature support for known relationships between the genes and the drug, as well as known interactions between the genes themselves.

### Summarized Benefits of Using Genvisage

By focusing on separating feature pairs, Genvisage offers researchers the ability to gain additional insight into their object classes beyond singular features, without the prolonged duration needed to train a complex machine learning model. By implementing optimizations that take advantage of a linear separability metric, Genvisage enables researchers to quickly explore their data, identify the strongest, most compelling features, and from simple visualizations form hypotheses about the interplay between features and with the object classes. The performance of our tool also allows researchers to investigate multiple definitions of the object classes and investigate alternative hypotheses interactively on the fly, as well as build a feature set to later pass to more in-depth, longer running machine learning-based analysis.

## 2 Methods

We begin by formally defining the *separability* problem, introducing our separability metric, and finally detailing optimizations that enable the rapid identification of the best separating feature pairs.

### 2.1 Problem Definition

Let 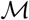 be a feature-object matrix of size *m* × *N*, where each row is a feature and each column is an object as shown in Figure 1. One example feature-object matrix is one where each object corresponds to a tissue sample from a cancer patient and each feature corresponds to a gene, where the (*i, j*)^*th*^ entry represents the expression level of the *i*^*th*^ gene in the *j*^*th*^ tissue sample. We denote the *m* features as 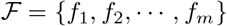 and *N* objects as *𝒪* = {*o*_1_, *o*_2_, …, *o*_*N*_}. Each entry 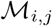 in 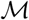 corresponds to the value of feature *f*_*i*_ for object *o*_*j*_ as illustrated in Figure 1.

**Fig 1.**
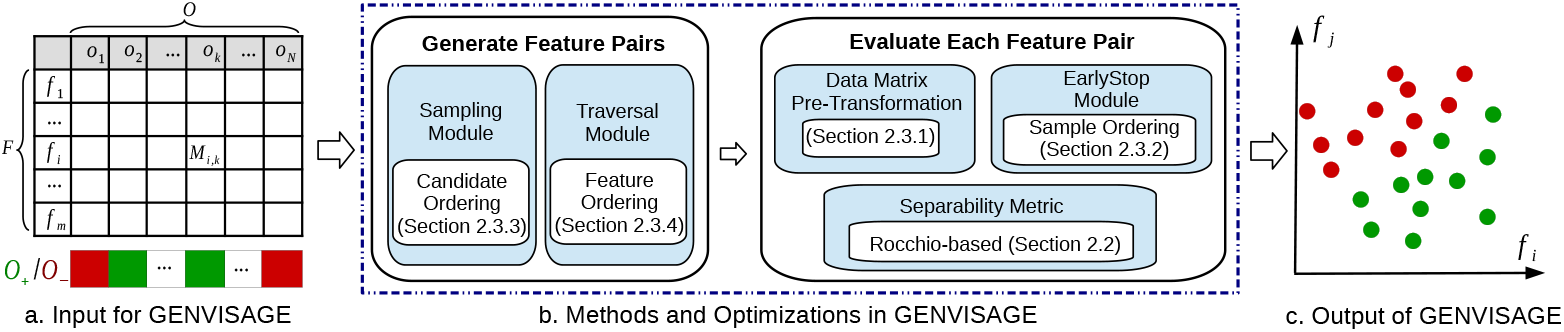
Genvisage Workflow. Given (left) a feature-object matrix and green positive and red negative class labels on the objects, Genvisage (center) evaluates all pairs of features using several optimizations to identify (right) the top feature pair and its corresponding visualization that best separates the object classes.

We are also given two non-overlapping sets of objects, one with a positive label, *𝒪*_+_ and the other with a negative label, *𝒪*_−_. In our example, tumor samples, *𝒪*_+_, may be assigned the positive label, and the healthy tissue samples, *𝒪*_−_, the negative label. The number of labeled objects, *n*, is equal to 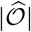 where 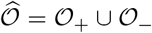. Also, let *l*_*k*_ be the label of object 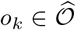, i.e., *l*_*k*_ = 1 if *o*_*k*_ is positive and *l*_*k*_ = −1 if *o*_*k*_ is negative.

Genvisage aims to find feature pairs that best separate the objects in *𝒪*_+_ from those in *𝒪*_−_ using only those features, and then output a visualization that demonstrates the separability (see Figure 1). (We will define the metric for separability subsequently.) A feature pair that leads to a good “visual” separation between the positive and the negative sets may be able to explain or characterize their differences via a interesting, non-trivial relationship among the features. The overall workflow is depicted in Figure 1. We now formally define the separability problem.

#### Problem 1 (Separability)

*Given a feature-object matrix 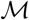 and two labeled object sets (*𝒪*_+_, *𝒪*_−_), identify the top-k feature pairs (*f*_*i*_, *f*_*j*_) that separate *𝒪*_+_ from *𝒪*_−_ based on a given separability metric*.

We will describe our separability metric in Section 2.2, and then discuss optimization techniques in Section 2.3. The notation used in the description of the method is summarized in Supplementary Table B.1.

### 2.2 Separability Metric

Given a feature pair (*f*_*i*_, *f*_*j*_) as axes, we can visualize the object sets *𝒪*_+_ and *𝒪*_−_ in a 2-D space, where each object corresponds to a point with x-value and y-value as the object’s value on feature *f*_*i*_ and *f*_*j*_ respectively. A desirable (i.e., both interesting and interpretable) visualization would be one in which the objects are *linearly separated*, defined as follows. Two sets of objects, i.e., *𝒪*_+_ and *𝒪*_−_, are said to be *linearly separable* [20] if there exists at least one straight line such that *𝒪*_+_ and *𝒪*_−_ are on opposite side of it. We focus on metrics that capture this linear separation, since it corresponds to an intuitive 2-D visualization. Given a feature pair (*f*_*i*_, *f*_*j*_) and a line *ℓ*, we can predict the label of an object *o*_*k*_, denoted as 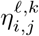, using Equation 1 below, where *w*_0_, *w*_*i*_ and *w*_*j*_ are coefficients of *ℓ* and *w*_*j*_ > 0:

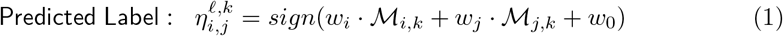

If *o*_*k*_ lies above the line *ℓ*, i.e., *o*_*k*_ has higher value on y-axis than the point on line *ℓ* with the same value on x-axis as *o*_*k*_, then 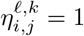; otherwise, 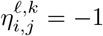. Let 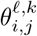 be the indicator variable denoting whether the sign of the predicted label matches the real label *l*_*k*_: if 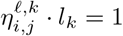, then 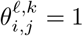; otherwise, 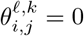.

Genvisage’s separability metric captures *how well the objects in the feature pair’s 2-D visualization can be linearly separated*, formally defined next. Given a feature pair (*f*_*i*_, *f*_*j*_) and a line *ℓ*, the separability score of the line (denoted 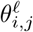) is defined as the sum of the indicators 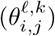 for all objects: 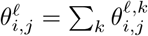. Figure 2(a) shows separability scores 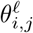 for different separating lines. For example, the separating line with 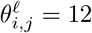 correctly separates six green points and six red points. The final separability score for a feature pair (*f*_*i*_, *f*_*j*_) (denoted as *θ*_*i,j*_) is defined as the best separability score 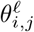 among all possible lines *ℓ*. Accordingly, we define the overall separability error of the feature pair as *err*_*i,j*_ = *n* − *θ*_*i,j*_.

**Fig 2.**
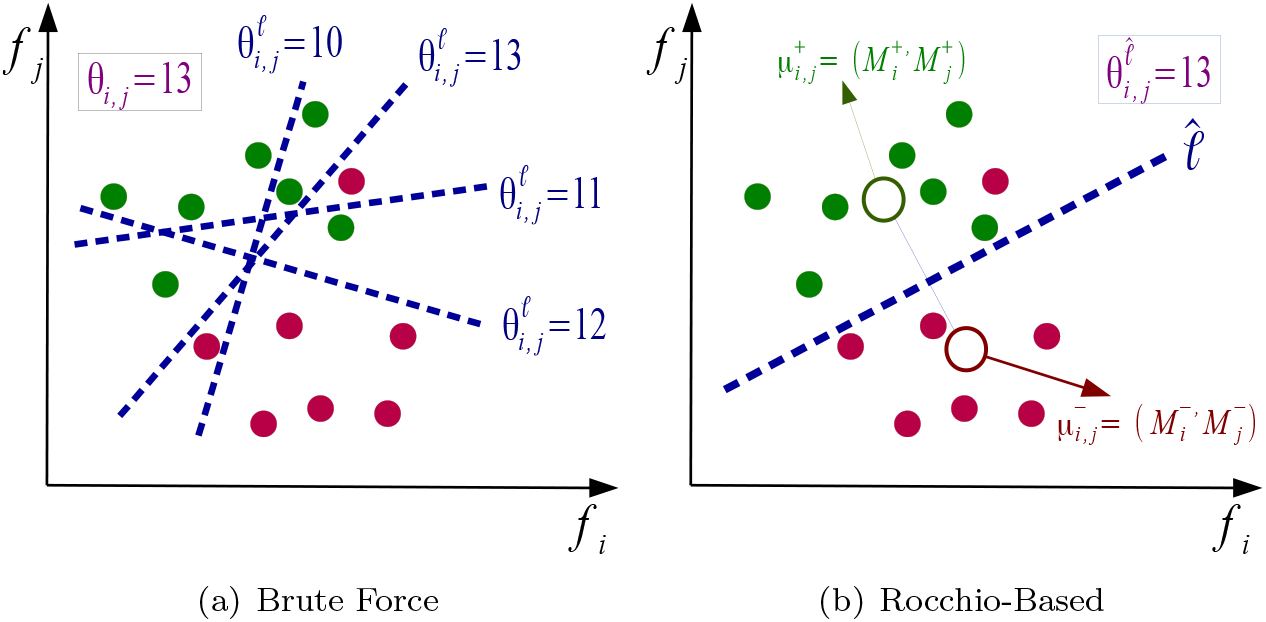
Calculating Separability Score *θ*_*i,j*_. The scored separating line can be defined using (a) brute force (few sample lines are shown) or (b) the representative line from a Rocchio-based measure based on the object class centroids (white circles).

#### Brute Force Calculation of *θ*_*i,j*_

As suggested in Figure 2(a), the simplest way to calculate *θ*_*i,j*_ is to first enumerate all possible separating lines *ℓ* and calculate 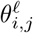 for each of them. We can easily trim down the search space to *O*(*n*^2^) lines by linking the points corresponding to every two objects in the 2-D plane. This is because the results of all other possible lines can be covered by these *O*(*n*^2^) lines [21]. Nevertheless, it is still very time-consuming to consider *O*(*n*^2^) lines for each feature pair (*f*_*i*_, *f*_*j*_).

#### Rocchio-based Measure

We can speed up the process by selecting a single *representative line L* providing us with an estimate of the true separability score *θ*_*i,j*_. In order to achieve a fast and reliable estimate, we select our representative line based on Rocchio’s algorithm [22]. Let us denote the centroids of positive objects *𝒪*_+_ and negative objects *𝒪*_−_ for a given (*f*_*i*_, *f*_*j*_) as 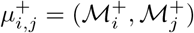 and 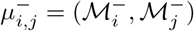 respectively, where 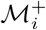 and 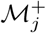 are the values of the centroids of the positive objects on feature *f*_*i*_ and *f*_*j*_, and 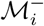 and 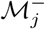 are the values of the centroids of the negative objects on feature *f*_*i*_ and *f*_*j*_. The perpendicular bisector of the line joining the two centroids is selected as the representative separating line *L* (see Figure 2(b)), with its coefficients corresponding to Equation 1 defined as 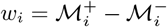, 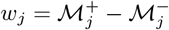, and 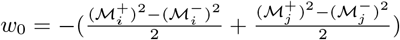.

#### Brute-force vs. Rocchio-based

Compared to the brute force calculation, the Rocchio-based measure is much more light-weight, but at the cost of accuracy in calculating *θ*_*i,j*_. Intuitively, the representative line is a reasonable proxy to the best separating line since the Rocchio-based measure assigns each object to its nearest centroid. We further empirically demonstrate that 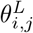 is a good proxy for *θ*_*i,j*_ in Section 3.2. Thus, we will focus on the Rocchio-based measure subsequently, removing *L* (or *ℓ*) from the superscripts where it appears, and using *θ*_*i,j*_ and 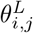 interchangeably.

### 2.3 Proposed Suite of Optimizations

In this section, we first analyze the time complexity of identifying the top-k feature pairs using the Rocchio-based measure, and then propose several optimization techniques to reduce the complexity.

#### Time Complexity Analysis

For a given feature pair (*f*_*i*_, *f*_*j*_), if we have already calculated the class centroids for each feature, the separating line *L* can be calculated in *O*(1). We can then calculate the number of correctly separated objects *θ*_*i,j*_ via *O*(*n*) evaluations. Since there are *O*(*m*^2^) feature pair candidates, the total time complexity is *O*(*m*^2^*n*), which can be very large, since *m* and *n* are typically large.

#### Optimizations: Overview

To reduce the time complexity, we introduce two categories of optimizations: those that reduce the amount of time for fully evaluating a given feature pair (Section 2.3.1, 2.3.2) and those that reduce the number of feature pairs that require full evaluation (Section 2.3.3, 2.3.4). In the following, we refer to these optimizations as *modules* to indicate that they can be used in any combination—however, in reality, careful engineering is necessary to “stitch” these modules together to multiply the effects of the optimizations.

Transformation module (Section 2.3.1) reduces redundant calculations across feature pairs by mapping the feature-object matrix 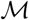 into a new space that enables faster evaluation of object labeling. EarlyStop module (Section 2.3.2) takes advantage of the fact that evaluation of a poorly separating feature pair can be terminated early without having to evaluate the separability of all *n* objects.

Sampling module (Section 2.3.3) first identifies likely top-k feature pair candidates by evaluating their separability on a sampled subset of all objects, and then conducts full evaluations only on these feature pair candidates. Finally, Traversal module (Section 2.3.4) reduces the number of feature pairs checked by greedily choosing feature pairs based on the separability of the corresponding single features. These optimization modules can be used on their own or combined with each other. In Section 3, we will show how these optimizations modules greatly reduce the running time of finding the top-k separating feature pairs without significantly affecting the accuracy.

##### 2.3.1 Pre-Transformation for Faster Feature Pair Evaluation

We observe that there is massive redundancy across *θ*_*i,j*_’s computation of different feature pairs. Motivated by this, we propose the Transformation optimization module which will pre-calculate some common computational components once across different features and reuse these components in evaluating the separability for each different feature pair. This Transformation module transforms the original 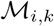 matrix into another space 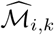 using the identified common feature pair components and updates the separability score equation accordingly. Specifically, with this transformation of the feature-object matrix 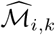, evaluating whether an object was correctly separated is simplified as: if 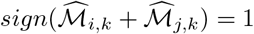, then 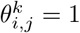; otherwise, 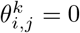. Details and an example can be found in Supplementary Note A.1 and Supplementary Figure C.1.

##### 2.3.2 Early Termination

>Given a feature pair (*f*_*i*_, *f*_*j*_), we need to scan all the objects to compute the separability score *θ*_*i,j*_. However, since we only need to identify feature pairs in the top-k, we can stop for each feature pair as soon as we can make that determination, without scanning all objects; we call this the EarlyStop module.

###### High Level Idea

We maintain a upper bound *τ* for the separability error *err*_*i,j*_ of the top-k feature pairs. Then, the lower bound of the separability score can be denoted as (*n − τ*). Given a feature pair (*f*_*i*_, *f*_*j*_), we start to scan the object list until the number of incorrectly classified objects exceeds *τ*. If so, we can terminate early and prune this feature pair since it cannot be among the top-k. Otherwise, (*f*_*i*_, *f*_*j*_) is added to the top-k feature pair set and we update *τ* accordingly.

###### Enhancement by Object Ordering

Although EearlyStop has the potential to always reduce the running time, its benefits are sensitive to the ordering of the objects for evaluation. Since we terminate as soon as we find *τ* incorrectly classified objects, we can improve our running time if we examine “problematic” objects that are unlikely to be correctly classified relatively early. For this, we order the objects in descending order of the number of single features *f*_*i*_ that incorrectly classify the object *o*_*k*_, i.e., 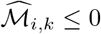. Thus, the first object evaluated is the one that is incorrectly classified by the most single features. The benefit of this strategy is illustrated with an example in Figure 3(a).

**Fig 3.**
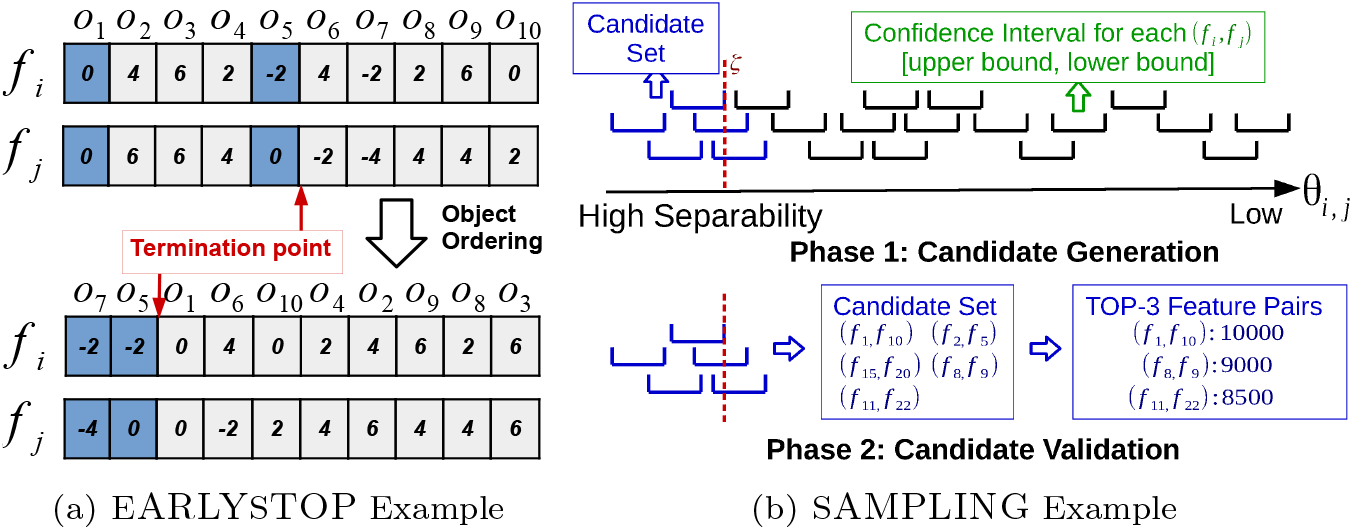
Optimization Module Examples. (a) When evaluating a feature pair with EarlyStop module, the transformed 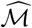 scores are scanned left to right and each incorrectly classified object is marked in blue. Without object ordering (above), evaluation terminates after five checked objects. When objects are reordered by the most “problematic” (below), the feature pair is rejected after checking only the first two objects. (b) To calculate the top-3 feature pairs with Sampling, the confidence interval of *θ*_*i,j*_ is calculated for every feature pair evaluated on the sample set 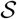 (above). The 3^*rd*^ interval lower bound *ζ* is obtained (red dotted line), and all feature pairs with a larger upper bound are designated as candidates for validation (blue intervals). The selected candidates (center box) are evaluated on the whole object set 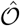 to compute the exact *θ*_*i,j*_ and pick the top-3 (right box).

##### 2.3.3 Sampling-based Estimation

One downside of the EearlyStop module is that the improvement in the running time is highly data-dependent. Here, we propose a stochastic method, called Sampling, that reduces the number of examined objects. Instead of calculating *θ*_*i,j*_ over the whole object set 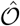, Sampling works on a sample set drawn from 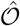.

###### High Level Idea

Sampling primarily consists of two phases: *candidate generation* and *validation* (Figure 3(b)). In phase one, we estimate the confidence interval of *θ*_*i,j*_ for each feature pair using a sampled set of objects and generate the candidate feature pairs for full evaluation based on where their confidence intervals lie. If the confidence interval overlaps with the score range of the current top-k, then it is selected for evaluation. In phase two (lower half of Figure 3(b)), we evaluate only the feature pairs in the candidate set, calculating *θ*_*i,j*_ over the whole object set, 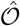, to obtain the final top-k feature pairs. Unlike our previous optimizations, Sampling returns an approximation of the top-k ranking feature pairs.

###### Candidate Generation

Let 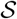 be a sample set drawn uniformly from 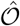. Given a feature pair (*f*_*i*_, *f*_*j*_), let 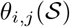 be the number of correctly separated objects in 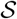. We can estimate 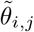 from 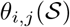 using 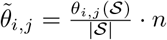 by assuming the ratio of correctly separated samples in 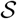 is the same as that in 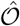. Using Hoeffding’s inequality [23], we have that by selecting 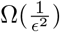 samples, that *θ*_*i,j*_ is in the confidence interval 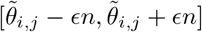 with high probability (details in Supplementary Note A.2). Since the sample size is 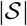 independent of the number of objects, this module helps Genvisage scale to datasets with large *n*.

Following the top half of Figure 3(b), we can first calculate the confidence interval of *θ*_*i,j*_ for each feature pair (*f*_*i*_, *f*_*j*_). Next, we compute the lower bound of *θ*_*i,j*_ for the top-k feature pairs, denoted as *ζ* as shown by the red dotted line. Finally, we can prune feature pairs away whose upper bound is smaller than *ζ*, keeping the candidate set *𝒞* of feature pairs depicted by blue intervals. These feature pairs *𝒞* will be further validated in phase two, i.e., candidate validation. Typically, |*𝒞*| will be orders of magnitude smaller than *m*^2^, the original search space for all feature pairs.

###### Candidate Validation

We re-evaluate all of the candidates generated from phase one to produce our final feature pair ranking. This evaluation is performed using the whole object set 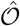 and the top-k feature pairs are reported (lower half of Figure 3(b)).

###### Enhancement by Candidate Ordering

In Section 2.3.2 we proposed an enhancement that allows us to terminate computation early by manipulating the order of the objects; here we similarly found a way to reduce the running time by changing the order in which feature pair candidates are validated in phase two. Instead of directly validating each feature pair candidate, we first order the candidates in descending order according to the upper bound of each candidate’s confidence interval. Then, we sequentially calculate the full separability score *θ*_*i,j*_ for each feature pair, and update *ζ* correspondingly. Recall that *ζ* is the current estimate of the lower bound of *θ*_*i,j*_ for the top-k feature pairs. Finally, we terminate our feature pair validation when the next feature pair’s upper bound smaller than the current value of *ζ* (Figure 4).

**Fig 4.**
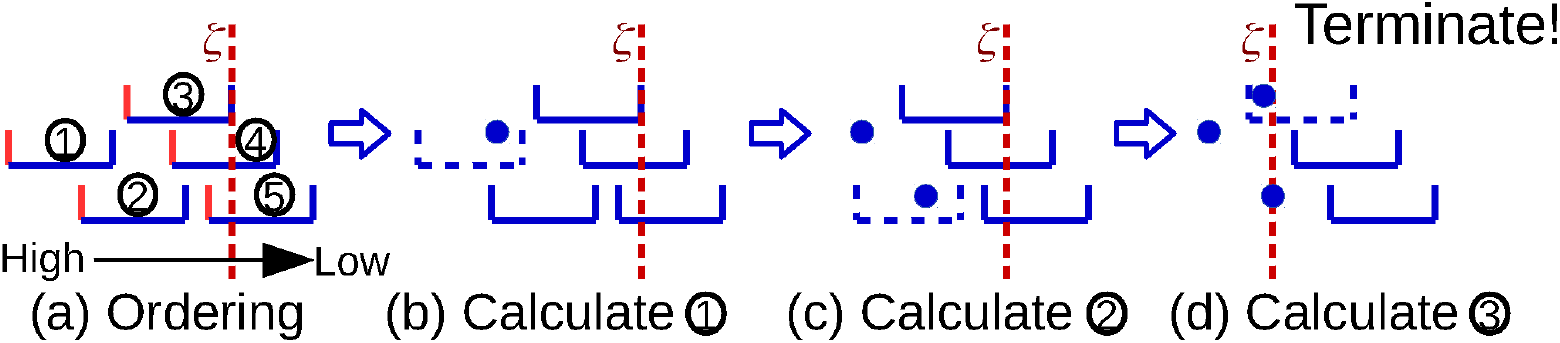
Candidate Ordering Enhancement. (a) Feature pair candidates are sorted by the upper bounds of their confidence intervals (solid red boundary), and the lower bound of the top-3 feature pairs, i.e., *ζ*, is set (red dotted line). (b,c,d) For each feature pair, we calculate *θ*_*i,j*_ (filled blue circle) using all objects and update *ζ* if necessary. Note that *ζ* is increased in (d) after evaluating the third feature pair and since *ζ* is larger than the upper bound of the fourth feature pair, candidate validation can terminate and return the top ranking pairs.

##### 2.3.4 Search Space Traversal

The optimizations discussed so far check fewer than *n* objects for each feature pair and reduce the number of feature pairs for full evaluation. Our Traversal module aims to reduce the number of feature pairs considered from *m*^2^ to a smaller number. Instead of examining each feature pair, we only examine a limited number of feature pairs, but in an optimized traversal order. The number of examined feature pairs, *χ*, determines a trade-off between efficiency and accuracy. Fewer feature pairs checked will result in faster running times, though at the cost of accuracy to the top-k. The order of the feature pairs must be determined carefully and we propose two alternative orderings based on the ranking of single features by their separability scores *θ_i,i_*. The first traversal order, called *horizontal traversal*, prioritizes feature pairs that have at least one high ranking single feature in the considered feature pair. The second order, called *vertical traversal*, prioritizes feature pairs where both features have high single feature scores rankings. See Supplementary Figure C.2 for more details and an example.

## 3 Results

In this section, we illustrate that Genvisage rapidly identifies meaningful, significant, and interesting separating feature pairs in real biological datasets. First, we describe the datasets and the algorithms used in our evaluation. Each algorithm that we evaluate represents a combination of optimization modules for ranking top-k feature pairs using our Rocchio-based measure—we report the running time and accuracy of the algorithms. Second, we compare the top-k feature pairs returned by Genvisage with the corresponding top-k single features, and examine their significance and support in existing publications. Last, we present some sample visualizations to illustrate the separability of the object classes.

### 3.1 Evaluation Setup

#### Datasets

We consider datasets from two biological applications (see Table 1): (a) in MSigDB, we find gene annotations such as pathways and biological processes that separate the differentially expressed genes from the undisturbed genes in specific cancer studies; (b) in LINCS, we find genes whose expression levels can distinguish experiments in which specific drug treatments were administered from others.

**Table 1.** Dataset Statistics. For each dataset, the number of features *m*, objects *N*, sample size 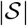 used by Sampling module, feature pairs *χ* examined by Traversal module, number of object sets: # of 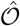, average positive set size: avg(|*𝒪*_+_|), and average negative set size: avg(*𝒪*_−_|).

| | $\vert \mathcal{F}\vert =m$ | $\vert \mathcal{O}\vert =N$ | $\vert \mathcal{S}\vert $ | $\chi$ | # of $\hat{\mathcal{O}}$ | avg($\vert \mathcal{O}_+\vert $) | avg($\vert \mathcal{O}_-\vert $) |
| --- | --- | --- | --- | --- | --- | --- | --- |
| MSigDB | 19,912 | 22,209 | 400 | $10^7$ | 10 | 295 | 21,914 |
| LINCS | 22,268 | 98,061 | 400 | $10^7$ | 40 | 165 | 97,897 |

In MSigDB, we are given a feature-object matrix with genes as the objects and gene properties as the features. Rather than being a 0/1 membership indicator matrix, the values of this feature-object matrix indicate the strength of the relationship between the gene and the set of genes that have been annotated with the gene property. Matrix values are calculated using random walks [24] on a heterogeneous network built from prior knowledge found in gene annotation and protein homology databases (see Supplementary Note A.3 for more details). The positive genes for each dataset in MSigDB are the set of differentially expressed genes (DEGs) in a specific cancer study downloaded from the Molecular Signatures Database (MSigDB) [25]. Each of our tests is an application of Genvisage to such a dataset, reporting pairs of properties that separate DEGs of that cancer study (the “positive” set) from all other genes (the “negative” set).

In LINCS, the feature-object matrix contains expression values for different genes (features) across many drug treatment experiments (objects) conducted on the MCF7 cell line by the LINCS L1000 project [26]. The values of the matrix are gene expression values as reported by the “level-4’ imputed z-scores measured in the L1000 project. In each dataset, the positive object set includes multiple experiments that used the same drug, at varying dosages and for varying durations. We applied Genvisage on each dataset so as to find the top pairs of genes (feature pairs) whose expression values separate the LINCS experiments relating to a single drug from all other LINCS experiments.

Note that the average number of positive objects in any dataset is far fewer than the average number of negative objects. To address this imbalance, we adjust 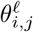 to a weighted sum form: 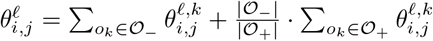.

##### Algorithms

We evaluated six combinations of our optimization modules from Section 2.3, listed in Table 2. For our baseline, we use the algorithm with only the matrix pre-transformation optimization module (Transformation). The rightmost column of Table 2 shows the varying time complexity of the algorithms. Consider the HorizSampOpt as an example. First, Transformation takes *O*(*mn*) time. Then, Traversal requires a sorting over the feature set, taking *O*(*m* log *m*) time. Finally, with Sampling over *χ* feature pairs, the running time is reduced from *O*(*m*^2^*n*) time to 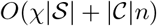 time, where the first and second term represent the time for candidate generation and candidate validation respectively. Note that |*𝒞*| is typically orders of magnitude smaller than *χ* in HorizSampOpt, as discussed in Section 2.3.3. Combinations of modules beyond the six reported were always inferior to one of the ones shown in the sense that they returned the same top-k feature pairs and had a longer running time. We implemented the algorithms in C++, and conducted the evaluations on a machine with 16 CPUs and 61.9 GB of RAM.

**Table 2.**
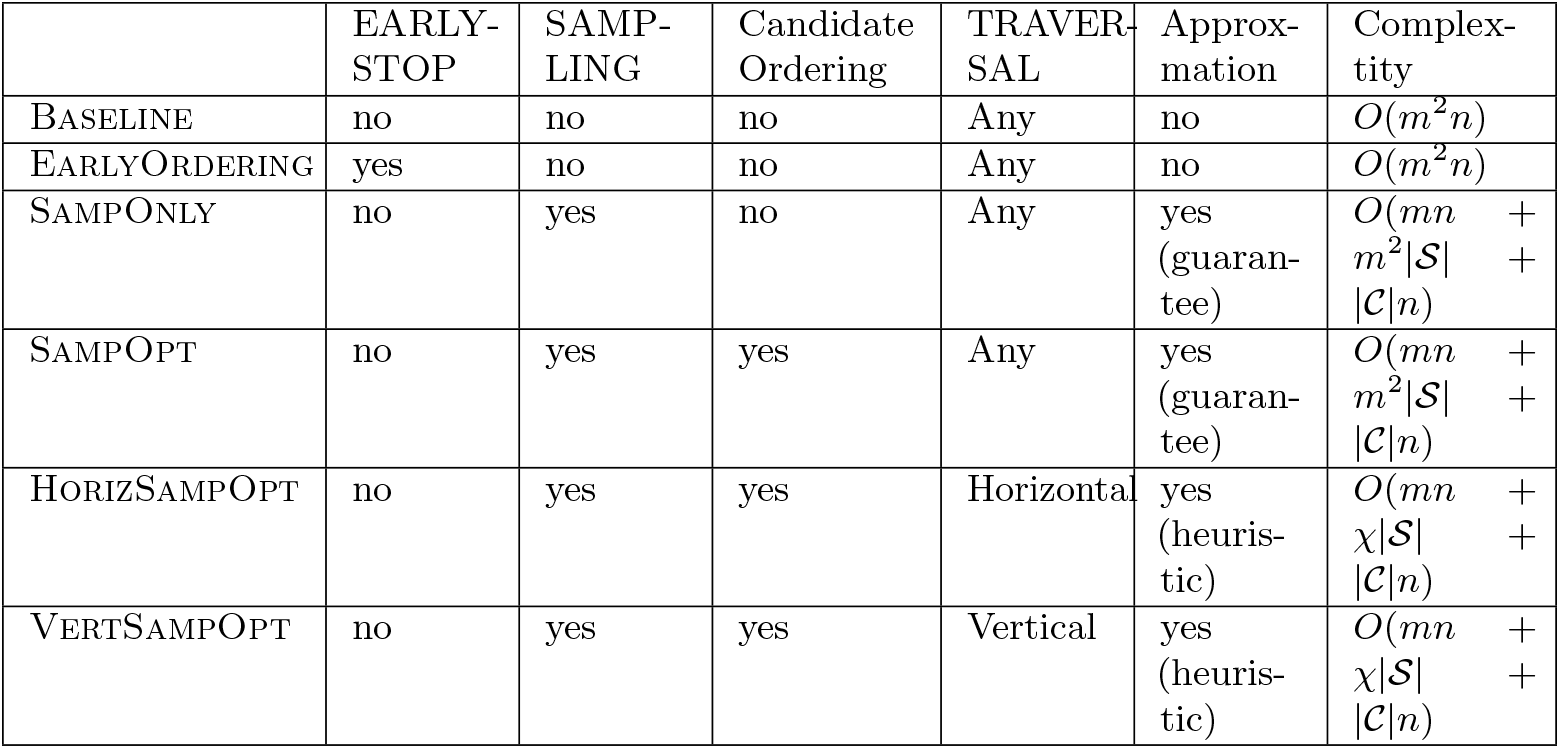
Selected Algorithms Using Different Optimization Modules. All algorithms, including the Baseline, are using Transformation. In addition, EearlyStop and Traversal are coupled with object ordering and feature ordering by default, respectively. For each algorithm (row), shows which optimization modules are employed, whether the algorithm is returning the exact answer or an approximation answer, and the running time complexity for that combination. The term “guarantee” (“heuristic”, resp.) indicates that the returned answer is with (without, resp.) stochastic guarantee. In addition, *m* and *n* are the number of features and objects, 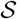 is the sampled set size, *χ* is the limit on the number of feature pairs considered, and *𝒞* is the number of generated feature pair candidates.

### 3.2 Comparison of Different Algorithms

In this section, we first justify that Rocchio-based measure is a good proxy for the best possible separating score computed by a brute force method. Then we compare the performance of the algorithms in terms of the running time and the separability of their top-1000 feature pairs.

#### Accuracy of Rocchio-based approximation

As discussed in Section 2.2, when using brute force, we need to consider *O*(*n*^2^) lines in order to find the best separating line 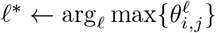, with a time complexity of *O*(*n*^2^*m*^2^) when considering all feature pairs. An alternative is to use Rocchio-based representative separating line *L*, dramatically reducing *O*(*n*^2^) lines considered to *O*(1). Since the brute force method becomes computationally infeasible for datasets with large *n*, we compared the Rocchio-based measure to the brute force-based measure using specially defined small object sets, 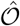, for the 10 datasets in MSigDB. For this comparison, the up-regulated genes in each MSigDB test was defined as the set of positive objects and the down-regulated genes as the set of negative objects, resulting in an average number of 295 objects for each comparison. We call the brute force-based separability score the *true* separability score, since it examines all possible separating lines. We first find the best feature pair using Rocchio-based measure and the brute force based measure separately (potentially different feature pairs) and then calculate the ratio of the true separability scores of the Rocchio versus the brute force best feature pairs. We observe that the Rocchio-based method picks a best feature pair that has true separability score similar to the best pair picked by brute force, with the ratio of the two scores being better than 0.94 in all ten datasets (Supplementary Figure C.3 (a)). Second, for the best feature pairs identified by Rocchio-based method for the ten datasets, we calculate the ratio of the Rocchio-based separability score and the brute force-based separability score, and find the difference to be greater than 0.96 on average (Supplementary Figure C.3 (b)).

#### Running Time

Figure 5 depicts the running times of our different selected algorithms. Each plotted box corresponds to one algorithm, representing the distribution of running times for finding the top-k feature pairs (by Rocchio score) for all datasets.

**Fig 5.**
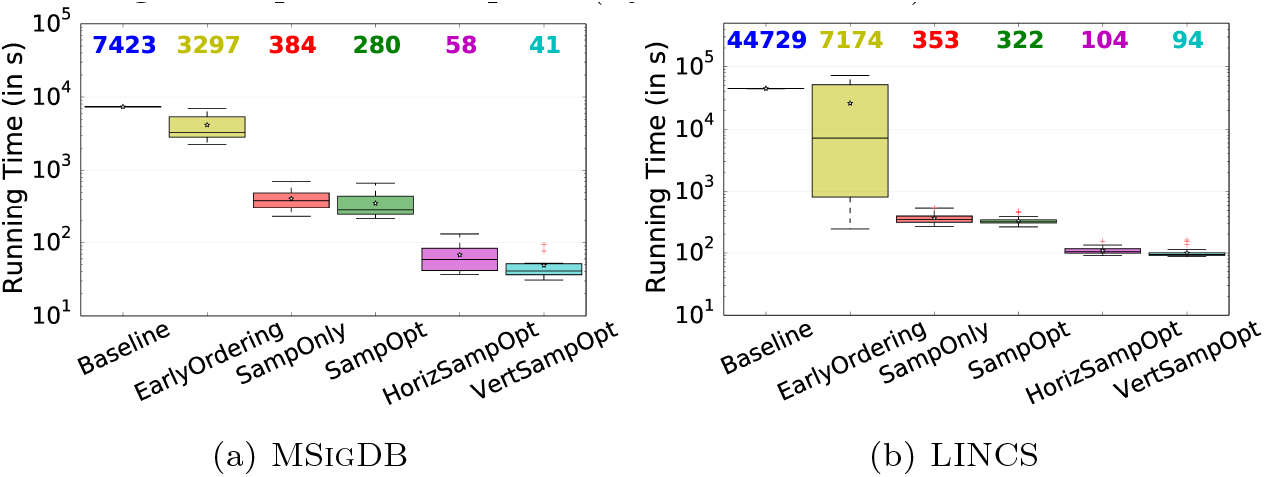
Running Time Comparison. A boxplot for each algorithm is shown with the median value appearing in matching color above. For each boxplot, whiskers are set to be 1.5× the interquartile range, the outliers are shows as red dots, and the average is marked with as a black star. The number on the top shows the median running time for each algorithm.

First, let us compare the median running times among different algorithms. For MSigDB, the Baseline takes more than 2 hours, EarlyOrdering takes less than 1 hour, SampOnly and SampOpt take around 6 and 5 minutes respectively, while HorizSampOpt and VertSampOpt both take only 1 minute on average. Overall, the optimizations result in a reduction of the running time by over 180×. We next examine the effect of different modules on the running time. *(a)* EearlyStop: we observe that the EearlyStop module helps achieve a 2× speed up, with the average number of checked objects (genes) reduced from 20*K* to 5*K* (Supplementary Table B.2); *(b)* Sampling: the Sampling module helps reduce the running time dramatically, with 20× reduction from Baseline to SampOpt, since on average only 2*M* candidates are generated from all possible 200*M* feature pairs (Supplementary Table B.2); *(c)* Traversal: the modules HorizSampOpt and VertSampOpt achieve an additional 6× speed-up compared to SampOpt by terminating after only considering *χ* = 10^7^ feature pairs, approximately 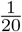 of all possible feature pairs. This speedup of HorizSampOpt and VertSampOpt is approaching the limit set by the feature ordering overhead (around 6*s*) and the time for the Transformation module (around 8*s*) (Supplementary Table B.2). The improvement over SampOpt is not stronger since the candidate generation phase of SampOpt is able to remove a vast amount of the feature pairs from full evaluation that would also be ignored by HorizSampOpt and VertSampOpt (Supplementary Table B.2).

Next, consider the log-scale interquartile range (IQR) of the running times for the different selected algorithms (Figure 5). We observe that EarlyOrdering has the largest interquartile range, indicating that the EearlyStop module, which tries to reduce the number of objects evaluated for each feature pair, is very dependent on the object set and feature values. As we discussed in Section 2.3.2, EearlyStop has no guarantee on improving the running time. In fact, the algorithm can occasionally be worse than the Baseline as shown in Figure 5(b) because EearlyStop incurs additional overhead for checking the criteria for pruning and early termination when scanning the object list for each feature pair. Similar results for LINCS are shown in Figure 5(b) (see Supplementary Note A.4).

#### Separability Quality

In Supplementary Figure C.3 (a), we found the the accuracy of the baseline method which computes the Rocchio-based estimate of top-k features to be high. The EearlyStop module is deterministic and produces the same top-k feature pairs as the baseline method only with optimized computation. The Sampling module, on the other hand, is stochastic and can only provide an approximation of the top-k feature pair ranking. Finally, the Traversal module is heuristic and may output top-k feature pairs that are very different from the ranking produced by the Baseline algorithm. and since Baseline returns the true Rocchio-based separability score of each feature pair, we measured the quality of each selected algorithm by counting the number of common feature pairs returned in the top-100 between the Baseline and the given algorithm. Figure 6 shows this separability quality comparison.

**Fig 6.**
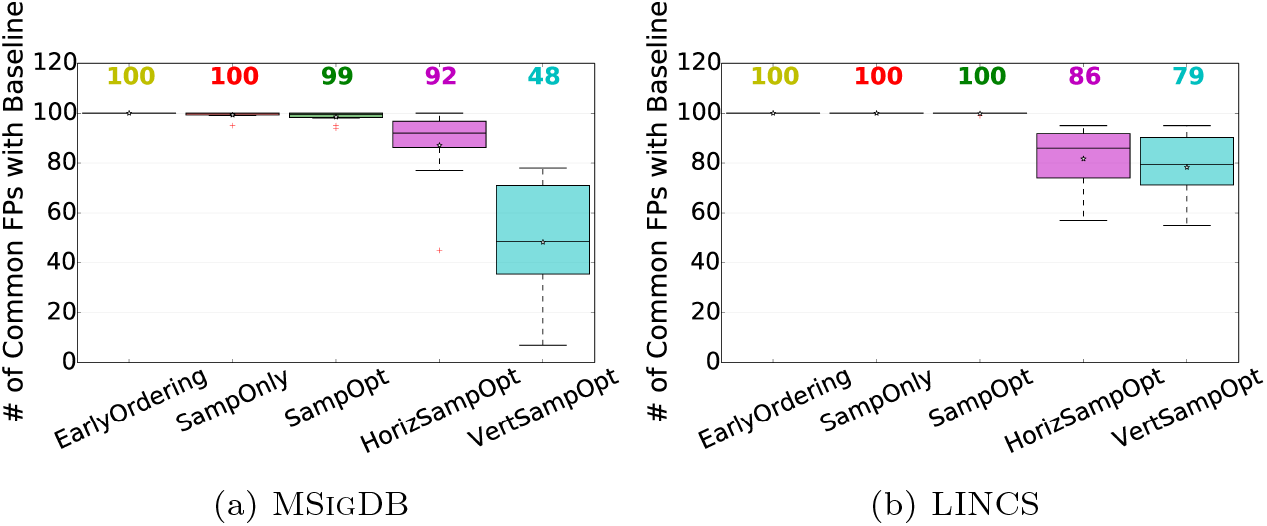
Separability Quality Comparison. Boxplots in the style of Figure 5 comparing the number of feature pairs each method returned from the 100 best feature pairs of the Baseline.

Let us first focus on MSigDB. EarlyOrdering, as expected, has exactly the same separability quality as the Baseline. We also observe that the SampOnly and SampOpt rankings are nearly identical to the top-100 feature pairs of the Baseline, owing to the probabilistic guarantee described in Supplementary Note A.2. The HorizSampOpt and VertSampOpt algorithms output a median of 92 and 48 feature pairs in common with Baseline, respectively, because of the heuristic Traversal module. In the MSigDB results, HorizSampOpt performs much better than VertSampOpt, with the median much higher and the interquartile range much narrower, as shown in Figure 6(a). This suggests, as we hypothesized, that interesting separating feature pairs exist outside of only the combinations of the top single features as in VertSampOpt. We repeated this quality analysis for LINCS and found that the Sampling based algorithms returned identical top-100 feature pairs for all 40 datasets. The quality of the Traversal based algorithms was again lower, though the performance separation of the HorizSampOpt and the VertSampOpt algorithms was not as large as for MSigDB.

#### Takeaways

If the accuracy is paramount, SampOpt is recommended; if the running time is paramount to the user, HorizSampOpt is recommended.

### 3.3 Feature Pair vs. Single Feature

In this section, we quantify the statistical significance of the top ranking results of the selected algorithms. We show that we often find separating feature pairs that are more significant than the best single separating feature. To assess the significance of a separating feature or feature pair, we first calculate the p-value using the one-sided Fisher’s exact test on a 2 × 2 contingency table. This contingency table is constructed with the rows being the true positive and negative labels, the columns being the predicted positive and negative labels, and the values being the number of objects that belong to each table cell. Using the Fisher’s exact test p-value, we assert that feature pairs can provide a better separability compared to single features, i.e., (a) feature pairs have stronger p-values compared to the corresponding individual features even after appropriate multiple hypothesis correction and (b) there exist high-ranked pairs of features that are poorly-ranked on their own as single features.

#### Single Feature

Finding top-k single features is a special case of finding feature pairs by setting *i* = *j*. For each single feature obtained, we compute the p-value with Fisher’s exact test, denoted as *pval*. Next, we define the Bonferroni corrected p-value as *corrected_pval* = *pval × m × n*, since there are *m × n* possible hypotheses, one for each possible single feature and separating line. We say a selected feature is significant if the corrected p-value is smaller than the threshold 10^*−*5^, i.e., −log_10_(*corrected_pval*) ≥ 5. In Figure 7, we plot the distribution of the corrected p-value of the top-100 features reported for each dataset in MSigDB and LINCS. We observe that 10 out of 10 datasets in MSigDB and 32 out of 40 datasets in LINCS have at least one significant single feature, and will focus on these datasets for further analysis. We observe very small p-values, ≤ 10^*−*50^, in the left part of Figure 7(a) and 7(b), indicating that single features are sufficient to separate the object classes for several datasets well.

**Fig 7.**
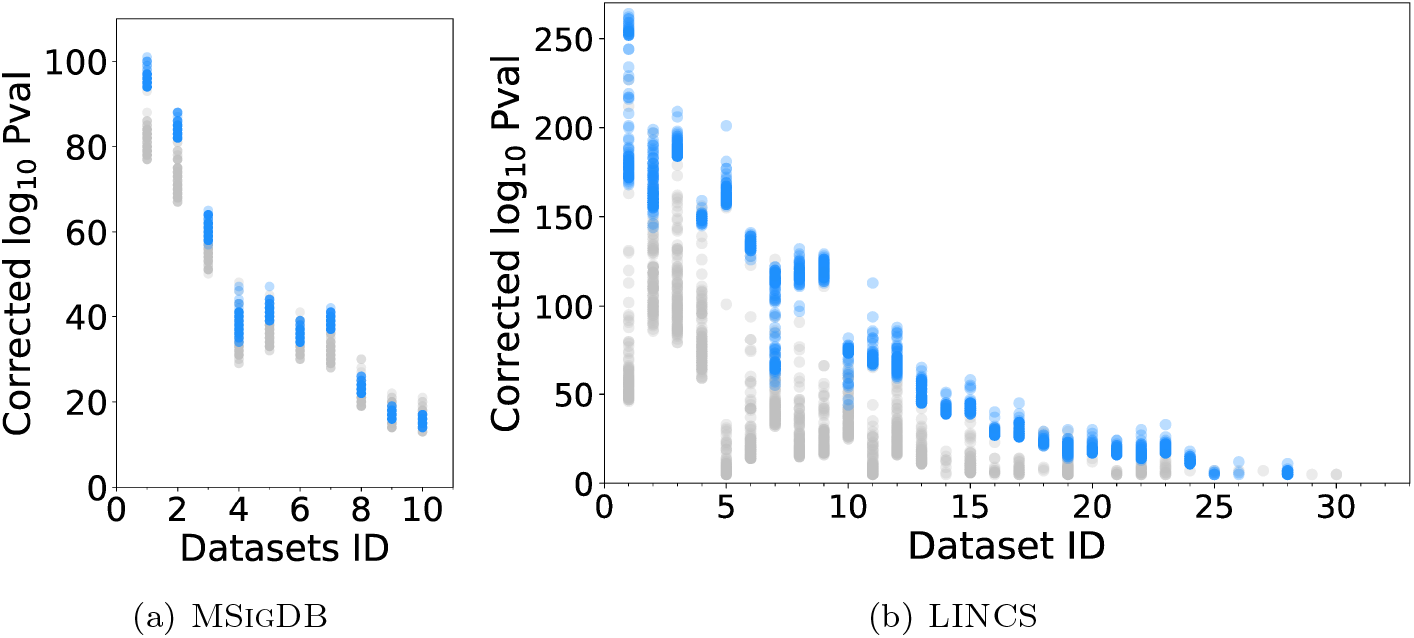
Single Feature Bonferroni Corrected P-value Distribution vs. Feature Pairs’ Corrected P-value Distribution. For each test (x-axis), shows the significance (−log_10_(*corrected_pval*)) of the top-100 best single features (grey dots) and feature pairs (blue dots) for the (a) MSigDB and (b) LINCS datasets. We order the datasets by their best corrected single feature p-value, and discard the datasets where no single feature has corrected p-value better than 10^*−*5^.

#### Feature Pair

We next build the contingency tables and calculate the p-value for the top-k feature pairs. To correct for *m*^2^ possible feature pairs and the *n*^2^ possible ways to choose the separating lines for each feature pair, we apply a Bonferroni p-value correction to produce the *corrected_pval* = *pval × m*^2^ × *n*^2^. We plot the distribution of the corrected p-values for the top-k feature pairs in Figure 7. Once again, the threshold for defining a significant feature pair is set to 10^*−*5^. We find that 10 out of 10 datasets in MSigDB and 27 out of selected 32 datasets in LINCS have at least one significant feature pair by this metric. Visual comparison of the top-100 single features to the top-100 feature pairs (Figure 7) per dataset reveals several datasets where the corrected p-values of the feature pairs are more significant than those of the best single features, even after accounting for the larger search space. Admittedly, this is not always the case, e.g., for five LINCS datasets no feature pair was found to be significant at *corrected_pval* ≤ 10^*−*5^ while at least one single feature did meet this threshold. Overall, this analysis suggests that rapid discovery of top feature pairs may identify more significant patterns in the given dataset than a traditional single-feature analysis does. In the following, we further illustrate that feature pairs can also provide better and newer insights compared to single features.

#### Improvement from Single Feature to Feature Pair

Having computed the corrected p-value for each single feature and feature pair in the top-100 for our datasets, we now examine the improvement of each feature pair from its two corresponding single features in terms of p-value. For each feature pair (*f*_*i*_, *f*_*j*_), we define the improvement quotient as the ratio between the corrected p-value of (*f*_*i*_, *f*_*j*_) and the better one of the corrected p-value of *f*_*i*_ or *f*_*j*_, i.e., 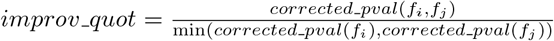. We examined only the *improv_quot* for the top-20 feature pairs for each of the 10 runs in MSigDB and 32 runs in LINCS. We found that on average across these datasets, 9.3 of the top-20 feature pairs in MSigDB and 8 of the top-20 feature pairs in LINCS are more significant than their corresponding single features (−log_10_(*improv_quot*) *>* 5). The distribution of the *improv_quot* is plotted in Supplementary Figure C.4. Overall, these histograms show that there is a improvement from single features to some feature pairs in terms of the separability significance. Next, we will explore the improved feature pairs more carefully, commenting on their redundancy, reliability, and relevance.

#### New Insights from Feature Pairs

In order to assess the quality of the top ranking feature pairs, we focused on the LINCS data set where the objects are experimental treatments on the MCF7 breast cancer cell line with the same *drug* and the features are expression values for different *genes*. For the evaluations above, we used object sets for the 40 drugs with the largest number of LINCS experiments. For the following analysis, we refine our list to those that are common drugs and have at least 60 LINCS experiments on the MCF7 cell line. These nine drugs are vorinostat, trichostatin, estradiol, tamoxifen, doxorubicin, gemcitabine, daunorubicin, idarubicin, and pravastatin. For each chosen drug, we ran the SampOpt algorithm of Genvisage to rank the top-1000 feature (gene) pairs for separating the LINCS experiments of the drug from all other MCF7 experiments.

For all drugs, except pravastatin, all of the top-1000 ranked feature pairs were found to be significant, i.e. −log_10_(*corrected_pval*) *>* 5 (see Table 3). As described in the Section 3.3, we are especially interested in feature pairs whose corrected p-value is better than the corrected p-values of their corresponding single features (−log_10_(*improv_quot*) *>* 0). We found 1070 “improved” feature pairs with larger separability over their single feature among the top1000 of these evaluation drug sets. One drug, trichostatin, had especially strong single features and showed no feature pairs that significantly improved on them. The remaining seven drugs, however, benefited from the feature pair analysis yielding between 9 (tamoxifen) and 369 (doxorubicin) improved feature pairs (Table 3).

**Table 3.** For each chosen drug from LINCS, the number of experiments in MCF7 cell line that were performed with that drug (NumExprs), and statistics for the top1000 feature pairs for that drug including the average *−* log_10_(*corrected_pval*) (Avg Signif), number of feature pairs with *−* log_10_(*corrected_pval*) *>* 5 (Top1000 Signif), and number with *−* log_10_(*improv_quot*) *>* 0 (Top1000 Improved).

| Drug | NumExprs | Avg Signif | Top1000 Signif | Top1000 Improved |
| --- | --- | --- | --- | --- |
| vorinostat | 904 | 235.5 | 1000 | 287 |
| trichostatin | 689 | 277.1 | 1000 | 0 |
| estradiol | 325 | 166.8 | 1000 | 203 |
| tamoxifen | 122 | 105.8 | 1000 | 9 |
| doxorubicin | 104 | 28.0 | 1000 | 369 |
| gemcitabine | 97 | 52.5 | 1000 | 116 |
| daunorubicin | 91 | 40.9 | 1000 | 28 |
| idarubicin | 78 | 30.1 | 1000 | 58 |
| pravastatin | 61 | -7.5 | 0 | 0 |
| Grand Total |  | 43.1 | 9068 | 1070 |

Many of the above-mentioned 1070 significantly improved feature pairs are partially redundant, in the sense that they comprise a common best-ranked single feature (gene). An example of this is with the object set for the drug (small molecule) estradiol. We found the gene PRSS23 as the single feature with the highest separability and many feature pairs containing PRSS23 and a second gene as having an improved corrected p-value, for example (PRSS23, RAP1GAP), (PRSS23, TSC22D3), and (PRSS23, BAMBI). We looked for evidence of the relationship between the drug estradiol and these feature pair genes in the Comparative Toxicogenomics Database (CTD) [27] and with our own literature survey. From this search, we found evidence for the pronounced effect of estradiol in increasing expression levels of PRSS23 [28], RAP1GAP [29], and BAMBI [30], and decreasing expression of TSC22D3 [31]. So although the top single feature (gene PRSS23) reoccurred in multiple top feature pairs, each secondary feature gene was also meaningfully related to the administered drug in this case.

We next examined the 1070 improved feature pairs, found over the 9 LINCS datasets, to determine their consistency with existing biological knowledge bases (see Supplementary Note A.5 for details). The interaction networks from these sources covered 23,167 genes and had at least one known interaction between 2.17% of all possible gene pairs. Of the 996 unique feature pairs with significant *improv_quot* where both genes mapped onto the genes covered by the interaction networks, 133 gene pairs (13.4%) were found to have at least one known interaction. This six-fold enrichment demonstrates that Genvisage more often finds pairs of genes that have a known relationship than is expected by chance. One example is (GLRX, NME7) that is especially good for separating vorinostat experiments from all others. Not only are both of these genes known to have increased mRNA expression in response to vorinostat [32], [33], but the two genes are annotated by STRING to both be in database pathways of nucleotide biosynthesis, co-express with each other in other model organisms, and mentioned together often in literature abstracts. Later, in Section 3.4, we will demonstrate that the positive objects and negative objects are visually separated under this feature pair, as in Figure 8.

**Fig 8.**
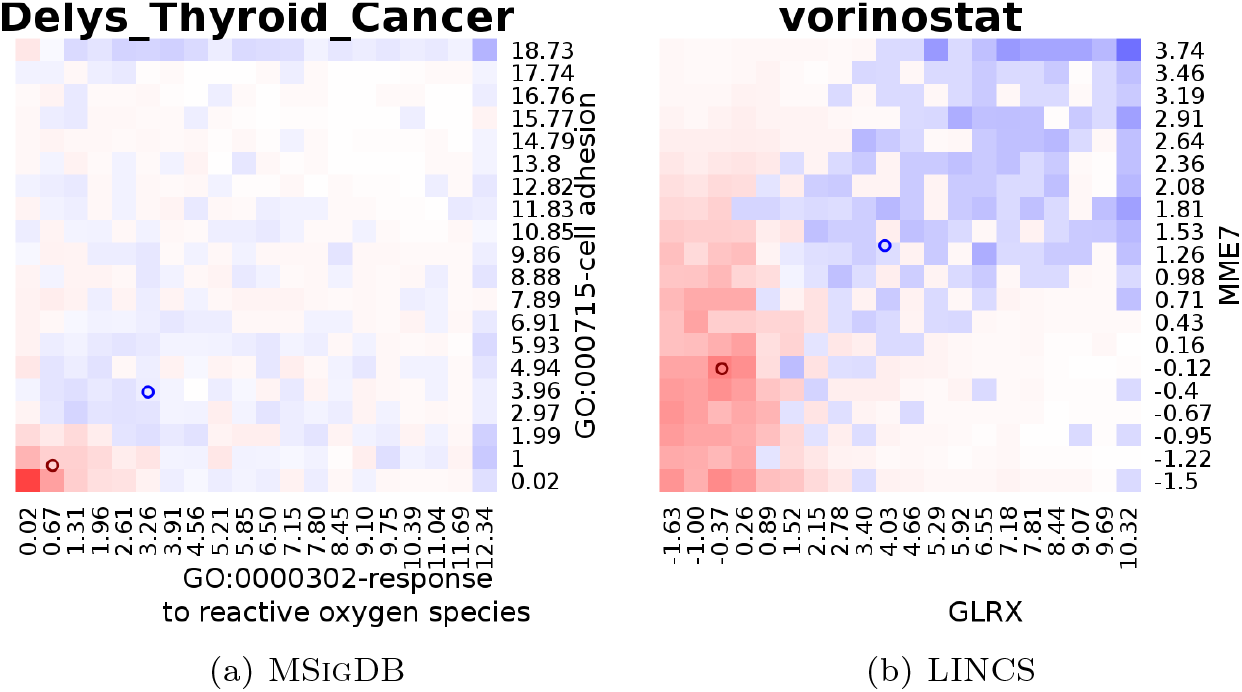
Visualization Output of Genvisage. Heatmap visualization with the pair of top features providing the x and y axes and the name of the run providing the plot title. The relative density of objects determines the color of the heatmap cells with blue indicating a greater proportion of positive objects and red indicating a greater proportion of negative objects. The class centroids are represented by blue (positive class) and red circles (negative class). The two examples shown are representatives from MSigDB and LINCS datasets.

In Supplementary Table B.3, we examine several of the “improved” feature gene pairs reported by Genvisage analysis for the LINCS nine drug datasets. Of thirty-nine feature pairs in this table, twelve of them have three types of accompanying evidence: 1) a literature-based relationship between the drug and the first gene, 2) a literature-based relationship between the drug and the second gene, and 3) an interaction network relationship between the pair of genes. Six have two of the three types of evidence and there are only three with no evidence at all. Particularly interesting are the top improved feature pairs in which neither of the single gene features ranked well alone. An example is the gene pair CDKN1A and CEBPB for separating doxorubicin experiments from others. Either gene feature alone is not within the top 600 genes for separating doxorubicin experiments from others. However, the combination of the pair is significant at a corrected p-value of 2 × 10^*−*25^ and is the second most improved feature pair for doxorubicin. This feature pair also has all three types of accompanying evidence; doxorubicin is known to increase expression of CDKN1A and CEBPB [34], and the pair of genes are annotated in STRING to have evidence for co-expression and text mining relationships. This feature pair can be used to form an interesting hypothesis for further analysis or experiment. The potential for finding more significant and previously unidentified features is why Genvisage is designed to recover top ranking feature pairs instead of just single features.

### 3.4 Output Visualizations

As discussed in Section 1, the output of Genvisage is not simply a ranking of the top feature pairs with their scores, but also a visualization that helps users to interpret the separability. In Figure 8, we depict sample output visualizations from the MSigDB and LINCS runs. For MSigDB, we select the feature pair with the highest improved p-value, i.e., *improv_quot*, using the SampOpt algorithm. For our LINCS representative, we visualize the gene feature pair (GLRX, NME7) for the drug vorinostat as described in the previous section. For the MSigDB example (Figure 8(a)), we observe that the feature values for negative objects are clustered around zero, while the genes differentially expressed in papillary thyroid carcinomas from this MSigDB study have larger values overall, indicating stronger connections to the two Gene Ontology terms features, cell adhesion and response to reactive oxygen species. This is consistent with studies that have highlighted the over expression of important cell adhesion genes in thyroid cancer [35]. For the LINCS example (Figure 8(b)), positive objects mostly have elevated expression for the two reported genes (GLRX and NME7) compared to the negative objects. The direction of this differential gene expression for both genes is consistent with literature for vorinostat experiments [32], [33]. These above two examples illustrate how visualization of significant feature pairs can be a useful way to explain the separability of object sets and understand the data.

## 4 Discussion

The Genvisage algorithm with its optimization modules enables researchers to visualize and explore the interplay between important pairs of genomic features rapidly, rather than relying on slow machine learning feature extraction methods or only examining the simple list of top single features. The optimization modules led to a two orders of magnitude speed up in the task of returning the top feature pairs for separating the biological classes in our two benchmark datasets, MSigDB and LINCS. The quality of these top feature pairs was confirmed by their agreement with literature and interaction databases, and the features are easily understood with intuitive heatmap visualizations. Genvisage relies on the Rocchio-based separability measure, which well approximates the best possible linear separator quickly and enables optimizations like Transformation that can pre-compute important quantities from the feature-object matrix before the positive and negative object sets are even provided. One potential downside of the Rocchio-based measure is that because of its dependency on linearity, feature pairs with distinct object class distributions that form complex, non-convex, non-isotropic patterns are potentially very interesting, but will not be well-ranked by Genvisage. Finally, in Genvisage, the optional Sampling module and Traversal modules make stochastic or greedy decisions in order to estimate the quality of and prune the potential candidate feature pairs for evaluation. While this greatly benefits the amount of time required to find the top ranking pairs, it has the potential to do so at the cost of ranking accuracy. Overall, we observed that for our settings, the sacrifice in accuracy was slight for the SampOpt feature pair rankings and more substantial when using the HorizSampOpt and VertSampOpt rankings with the greedy candidate traversal. However, users of Genvisage are able to optimize the trade-off with performance and accuracy by modifying the sample size, 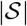, used by the Sampling module or the number of candidate feature pairs examined, *χ*, by Traversal module depending on the needs of their research and dataset.

## Appendix A Supplementary Notes

### A.1 Pre-Transformation Module

Let us review the process of computing the separability *θ*_*i,j*_. Given a feature pair (*f*_*i*_, *f*_*j*_) and the corresponding positive and negative centroids, *(i)* we first compute *w*_0_, *w*_*i*_ and *w*_*j*_ for *L*. Next, for each object *o*_*k*_, *(ii)* we obtain the predicted label 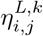 according to Equation 1. This step requires two multiplications and three additions. Finally, *(iii)* we calculate 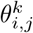 and the separability *θ*_*i,j*_ based on formulations in Section 2.2. This whole process is repeated for every feature pair candidate. However, there is massive redundancy across the processing of different feature pairs. For instance, when calculating the separability for two different feature pairs (*f*_*i*_, *f*_*j*_) and (*f*_*i*_, *f*_*j*′_) with a common *f*_*i*_, *w*_*i*_ is in fact shared, and calculation of 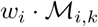 in Equation 1 is repeated for each object *o*_*k*_.

Given this, we propose the Transformation optimization module which will pre-calculate some common computational components once across different features and reuse these components the separability for each feature pair to eliminate the repeated computation. This Transformation module transforms the original 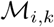 matrix into another space using our identified common feature pair components and updates the separability score equation accordingly.

For each feature *f*_*i*_, we find the average values of the positive and negative objects for that feature, 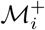 and 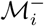 respectively, and then we pre-transform 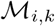, i.e., the value of object *o*_*k*_ on the feature *i*, to 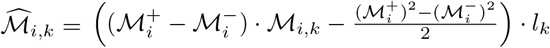. The basic idea is to decompose Equation 1 into two components, with each one only related to a single feature. This transformation incorporates the class centroids into the matrix values, obviating their integration later for every feature pair that involves the given feature. We also multiplies in the class label of the object, *l*_*k*_, rather than repeating this multiplication every time the object is evaluated (see example in Supplementary Figure C.1). With this transformation of the feature-object matrix, evaluating whether an object was correctly separated is simplified as: if 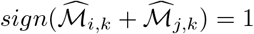, then 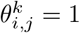; otherwise, 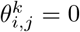. Note that this step only involves one addition and one comparison and is performed only once for each feature. Next, we can calculate overall separability score 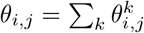. Overall, we not only eliminate the steps of computing *w*_0_, *w*_*i*_ and *w*_*j*_ for every feature pair, but also reduce the cost of calculating 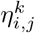 in Equation 1. With the Transformation module, we calculate 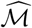 as a pre-transformation step, and use it when evaluating feature pairs instead of 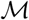 and calculate *θ*_*i,j*_ accordingly.

### A.2 Estimation Accuracy in Sampling Module

We have proposed Sampling for estimating *θ*_*i,j*_ in Section 2.3.3. Next, we formally quantize the sample set size in Theorem 1.

#### Theorem 1 (Estimation Accuracy)

*By considering* 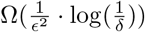 *samples, with probability at least* 1 − *δ, we have* 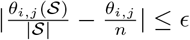, i.e., 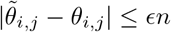.

We can treat log(1*/δ*) as a constant, e.g., by setting *δ* = 0.05. Thus, Theorem 1 essentially states that with only 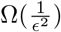 samples, with probability 95%, the confidence interval for *θ*_*i,j*_ is 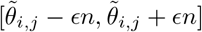.

### A.3 Construction of MSigDB Feature-Object Matrix

To construct the feature-object matrix for MSigDB, which has gene objects and gene property features, we collected prior knowledge about gene annotations and protein homology from several databases. Our gene annotations and gene properties were extracted from Gene Ontology terms [36], PFam domains [37], and Reactome [38] and KEGG [39] pathways. We constructed a heterogeneous network with nodes for all 22,210 genes and 21,235 properties from these databases and with edges representing their annotations between genes and properties. We also created weighted homology-based edges in the network between pairs of genes based on their protein sequence similarity as determined by BLAST scores [40]. We used the first phase of the DRaWR algorithm [24] with a restart probability of 0.5 to perform a random walk restarting from each gene node on the heterogeneous network, thereby scoring the connectivity of all nodes in the network to the gene. For each gene-property pair (*g*, *r*), we assigned the numeric value from the random walk stationary probability distribution that represents not only whether the gene is annotated with that property, but also whether other genes closely related to gene *g* are annotated with property *r*. We thus obtained a feature-object matrix describing each gene (object) as a vector of its strength of association with each property (feature) in light of prior biological knowledge.

### A.4 Speedup Analysis for LINCS

In Figure 5(b), we observe over 400× average decrease in the running time of finding the top-k feature pairs that separate the LINCS experiments of a single perturbagen from others. The greatest speedup comes with adding the Sampling module, where only 100*K* feature pair candidates, i.e.,|*𝒩*|, are checked out of all 250*M* feature pairs (Supplementary Table B.2). For the selected algorithms with best running times, HorizSampOpt and VertSampOpt, the pre-transformation and feature ordering overhead account for an average of 45 + 35 = 80*s* of the overall 104 and 94 median seconds respectively.

### A.5 Gene Interaction Datasets

For our knowledge bases of protein and gene interactions, we downloaded datasets derived from 8 data sources: STRING [41], Reactome [42], Pathway Commons [43], HumanNet [44], BioGRID [45], Intact [46], DIP [47], and BLAST [40] databases. The datasets were downloaded, harmonized, and mapped to Ensembl gene identifiers using the KnowEnG Knowledge Network Builder https://github.com/KnowEnG/KN_Builder. The final processed network used in this work can be downloaded from https://github.com/KnowEnG/KN_Fetcher/blob/master/Contents.md#gene.

## Appendix B Supplementary Tables

### B.1 Genvisage Method Notation

Notation used in this paper.

| Symb. | Description | Symb. | Description |
| --- | --- | --- | --- |
| $\mathcal{M}$ | feature-object matrix | $\mathcal{F}$ | feature set in $\mathcal{M}$ |
| $f_i$ | feature $i$ in $\mathcal{F}$ | $m$ | number of features in $\mathcal{F}$ |
| $\mathcal{O}$ | object set in $\mathcal{M}$ | $N$ | number of objects in $\mathcal{O}$ |
| $\mathcal{O}_+$ | positive object set | $\mathcal{O}_-$ | negative object set |
| $\hat{\mathcal{O}}$ | labelled object set | $n$ | number of labelled objects in $\hat{\mathcal{O}}$ |
| $o_k$ | object $k$ in $\hat{\mathcal{O}}$ | $l_k$ | label of object $o_k$ |
| $\ell$ | separating line in 2-D | $L$ | representative line in 2-D |
| $\eta_{i,j}^{\ell,k}$ | predicted label of $o_k$ | $\theta_{i,j}^{\ell,k}$ | $o_k$ is correctly separated? |
| $\theta_{i,j}^{\ell}$ | # correctly separated $o_k$ | $\theta_{i,j}$ | separability score |
| $\hat{\mathcal{M}}$ | $\mathcal{M}$ after transformation | $\hat{\theta}_{i,j}$ | estimated $\theta_{i,j}$ |

### B.2 Running Time Detailed Comparison

For each dataset collection, MSigDB and LINCS, show two statistics FPsChecked and ObjectsChecked, for each optimization module phase (columns). FPsChecked is the number of feature pairs evaluated in the phase, and ObjectsChecked is the average number of sample objects that are evaluated across all feature pairs.

### B.3 In-depth Analysis of ’Improved’ Feature Pairs

For each drug dataset (col C), we return a limited number of gene feature pairs (cols E,F) that after correction were the most ”improved” over their corresponding single feature results either by the change in the corrected pvalue (cols H,I,J,N,O) or the change in the feature rankings (cols K,L,M,P). Pubmed IDs (cols Q,R) are provided when the relationship between the drug and the gene feature were found in the Comparative Toxicogenomics Database (CTD) or by manual literature search (denoted by *-asterisk). When relationships between the gene features themselves were discovered, the type and strength of the relationships were reported (col S) and the number of relationships quantified (col T).

## Appendix C Supplementary Figures

### C.1 Example for Transformation Module

The Transformation module is applied once to the original values in the feature-object matrix 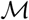 (above) to produce 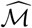 (below). For two features, *f*_*i*_ and *f*_*j*_. The top half depicts 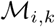 and 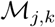 before transformation, where green color represents a positive label and red color represents a negative label. In this example, the centroids of the positive and negative objects are 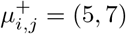 and 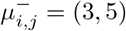 respectively. Hence, we can rewrite 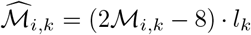 and 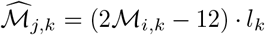 for features *f*_*i*_ and *f*_*j*_ respectively. After calculation, we can obtain the values for 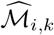 and 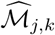 shown in the bottom half.

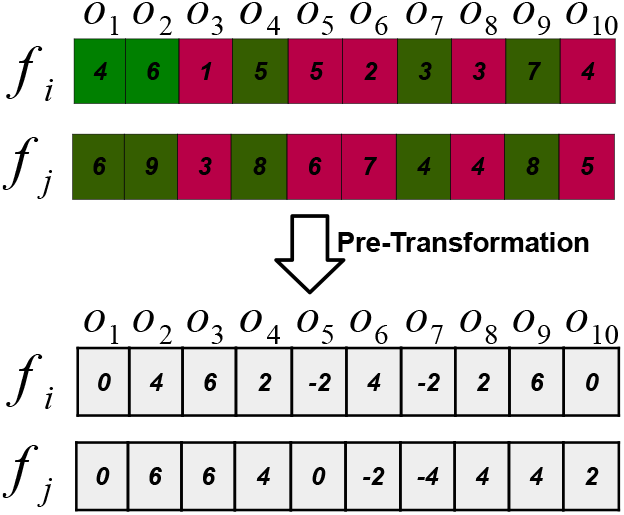

### C.2 Illustration for Traversal Module

We rank individual features based on their single feature separability scores, *θ*_*i,i*_, from best to worst, 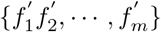.

- *Horizontal traversal:* For each feature 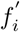, pair it with each other feature 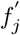, where *j ≥ i*, to obtain 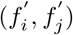. Repeat for each 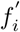, where 1 *≤ i ≤ m*.
- *Vertical traversal:* For each feature 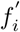, pair it with each other feature 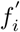, where *i ≤ j*, to obtain 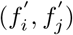. Repeat for each 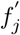, where 1 *≤ j ≤ m*.

For an example, suppose there are 20,000 features, *m* = 2 *×* 10^4^. Initially, the number of possible feature pairs is roughly 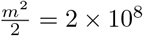. However, if we limit the number of considered feature pairs to *χ* = 10^7^, we reduce our search space to 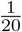 of the total number of feature pairs. We order the single features by their individual separability scores. In horizontal traversal, only feature pairs with at least one individual feature ranked in the top 500 will be considered; while vertical traversal will consider only feature pairs with both individual features ranked better than 2000.

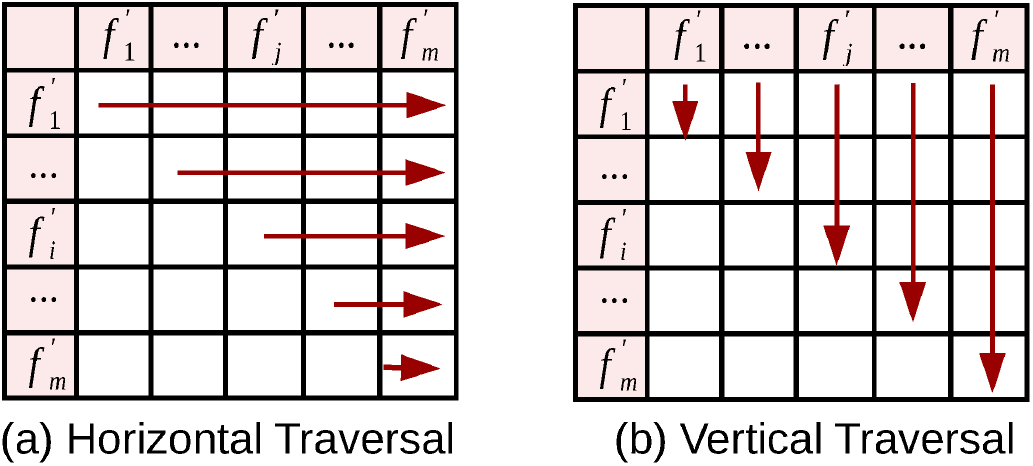

### C.3 Separability Score Comparison

Comparison of Brute Force-based and Rocchio-based separability score. (a) For each of 10 datasets, we display the ratio of the true separability score between the best feature pair chosen by brute force and by the Rocchio-based method. (b) For each dataset, we display the ratio of the true separability score and the Rocchio-based separability score for the best feature pair selected using Rocchio-based method.

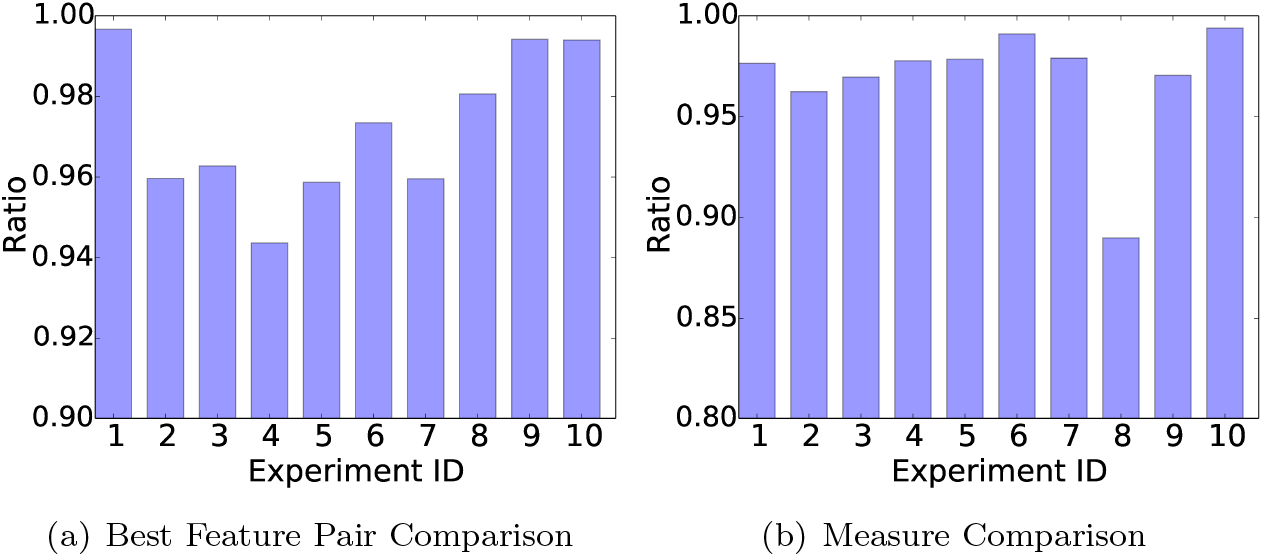

### C.4 Histogram of *improv_quot*

Histogram of *improv_quot*. For the top-20 feature pairs from all runs from the (a) MSigDB and (b) LINCS datasets, distribution of the improvement of the feature pair significance over the corresponding single feature significance. The red line shows the significance threshold of 5.

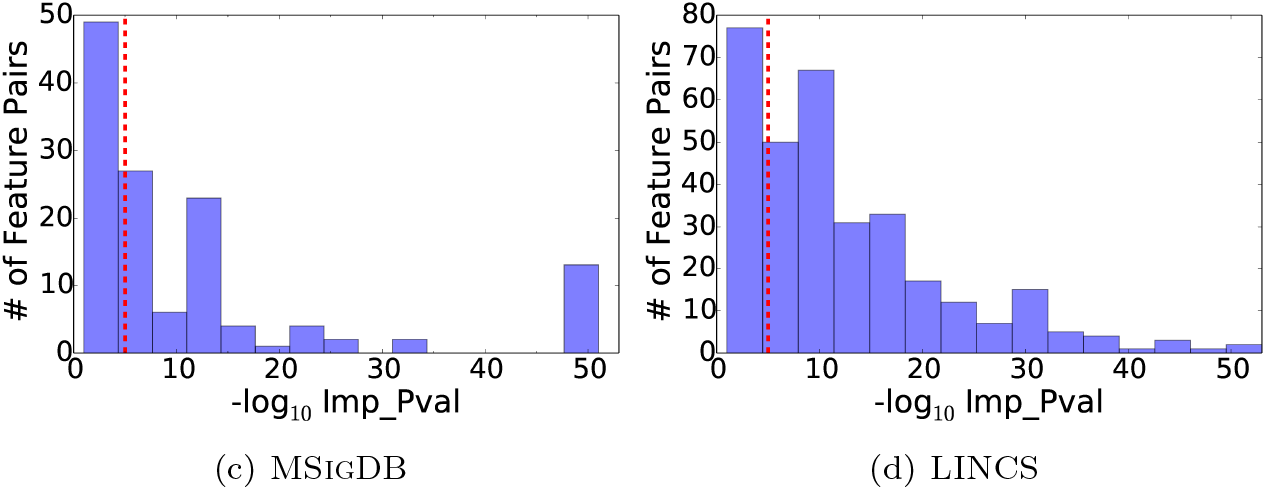

